# Multiscale interactome analysis coupled with off-target drug predictions reveals drug repurposing candidates for human coronavirus disease

**DOI:** 10.1101/2021.04.13.439274

**Authors:** Michael G. Sugiyama, Haotian Cui, Dar’ya S. Redka, Mehran Karimzadeh, Edurne Rujas, Hassaan Maan, Sikander Hayat, Kyle Cheung, Rahul Misra, Joseph B. McPhee, Russell D. Viirre, Andrew Haller, Roberto J. Botelho, Raffi Karshafian, Sarah A. Sabatinos, Gregory D. Fairn, Seyed Ali Madani Tonekaboni, Andreas Windemuth, Jean-Philippe Julien, Vijay Shahani, Stephen S. MacKinnon, Bo Wang, Costin N. Antonescu

## Abstract

The COVID-19 pandemic has led to an urgent need for the identification of new antiviral drug therapies that can be rapidly deployed to treat patients with this disease. COVID-19 is caused by infection with the human coronavirus SARS-CoV-2. We developed a computational approach to identify new antiviral drug targets and repurpose clinically-relevant drug compounds for the treatment of COVID-19. Our approach is based on graph convolutional networks (GCN) and involves multiscale host-virus interactome analysis coupled to off-target drug predictions. Cell-based experimental assessment reveals several clinically-relevant repurposing drug candidates predicted by the *in silico* analyses to have antiviral activity against human coronavirus infection. In particular, we identify the MET inhibitor capmatinib as having potent and broad antiviral activity against several coronaviruses in a MET-independent manner, as well as novel roles for host cell proteins such as IRAK1/4 in supporting human coronavirus infection, which can inform further drug discovery studies.

## Introduction

Human coronaviruses have emerged as significant agents in serious human disease, highlighting the need for rapid development of therapeutics. Recently, the severe acute respiratory syndrome (SARS) coronavirus-2 (SARS-CoV-2) has caused a pandemic with >125M individuals infected and over 2.5M deaths globally ^1^. Prior to this, the related human coronaviruses SARS-CoV-1 and Middle East Respiratory Syndrome coronavirus (MERS-CoV) were responsible for outbreaks of severe human coronavirus disease with significant mortality ^2,3^. Additional human coronaviruses including NL63, OC43, and 229E elicit milder disease such as the common cold ^4^, although more severe human diseases related to these viruses have been reported ^5^. Human coronaviruses thus represent an important family of related viruses that impact human health worldwide with new therapies for these agents urgently needed. The recent emergence of SARS-CoV-2 variants of concern ^6,7^ indicates that therapeutic strategies with broad antiviral activity to human coronaviruses are a high priority.

SARS-CoV-2, SARS-CoV-1 and NL63 enter cells by using their Spike (S) outer protein, which interacts with angiotensin converting enzyme 2 (ACE2) on host cells ^8–11^, while 229E and OC43 depend on host cell aminopeptidase N and glycosaminoglycans, respectively ^12,13^. In addition, other host cell receptors contribute to SARS-CoV-2 entry, such as neurolipin-1 ^14,15^. The coronavirus-bound receptor protein(s) enter cells via clathrin-mediated endocytosis ^16,17^ or other membrane traffic mechanisms ^18^, after which the viral RNA genome undergoes replication and expression of viral proteins, leading to assembly of viral progeny at a specialized ER-derived coronavirus replication organelle ^19^. This is followed by coronavirus release to the extracellular milieu by a mechanism involving lysosomal exocytosis ^20^, allowing spread of the viral particles to nearby cells ^21,22^.

Human coronavirus cellular entry, replication, assembly and egress depends on a wide range of host cell proteins and functions. For SARS-CoV-2, 26 viral proteins were found to interact with 332 host cell proteins, spanning a range of functions including membrane trafficking, centrosome structure and function, membrane transport, DNA replication and stress granule formation ^23^. This viral protein interactome provides a rich source of information from which to identify host proteins that are functionally important for viral entry and replication and thus may serve as antiviral drug targets. Infection studies have also demonstrated conserved host-protein interaction patterns across different coronaviruses, suggesting that some host proteins can be targeted for broader antiviral activities ^24–26^. The virus/host protein interactome and the identification of proteins functionally required for viral entry and replication of common cold coronaviruses may reveal important novel pan-human coronavirus antiviral drug targets.

Antivirals targeting SARS-CoV-2 such as remdesivir have modest clinical benefits ^27^, indicating a need for more effective antivirals to complement anti-inflammatory therapy approaches for the treatment of COVID-19. As the timeline for development of new drugs is ~10 years ^28^, and given the current dire need for effective antiviral agents, the identification of new drugs with antiviral activity cannot provide antiviral therapies in time to address the current pandemic. Drug repurposing is a preferred method for developing rapid response therapies ^28,29^, as it prioritizes the identification of additional drugs that can be repurposed as antivirals. Successful drug repurposing discoveries to date have been largely accidental or hypothesis-driven ^30^.

Computational approaches provide an opportunity to identify repurposing candidates that may have been overlooked by hypothesis-driven approaches or missed by less robust high-throughput screening approaches. These hypothesis-free strategies reference broad data sources to identify new protein targets, identify new compounds for a pre-selected target, or pair phenotypic signatures of a disease state with drug actions ^31^. Notably, multiscale interactome approaches combine known relationships between disease, biological pathways, genes, proteins, and other - omic data to predict new potential indirect relationships ^32^. In biological systems however, the correlation between the physical biological interactions and the probability of possible treatment is hidden, representing a major challenge when ranking possible indirect relationships. Graph Convolutional Networks (GCNs) offer a systematic, Machine-Learning (ML) based approach to model the value of each relationship in the network empirically based on their intrinsic properties ^33,34^

Drug-target coverage in the human proteome is another major challenge to drug repurposing. As of 2019, only 667 of the roughly 20,000 human proteins (~3%) are directly targeted by FDA-approved drugs ^35^. *In silico* virtual screening approaches can provide interaction predictions for proteins that are not established drug targets, but otherwise have high disease relevance. ML-based virtual screening approaches are capable of cross-screening libraries of clinically relevant compounds with large sets of proteins, to assist repurposing programs with multiple plausible targets ^36^.

In this study, we combine GCN-based multiscale host-viral interactome approaches for target discovery with off-target interaction predictions from the PolypharmDB database ^36^ to shortlist clinically-relevant repurposing candidates for screening in coronavirus infectivity assays. We then examine the effectiveness of these predictions using a combination of human coronavirus entry and infection assays. We reveal several compounds that have antiviral activity, in particular capmatinib, a drug known to inhibit the receptor tyrosine kinase MET. Notably, we find that capmatinib has potent anti-human coronavirus activity in a MET-independent manner. Further, we find novel roles for human proteins such as IRAK1/4 in supporting human coronavirus infection.

## Results

A total of 26 drug repurposing candidates for SARS-CoV-2 were identified using three separate approaches involving a multiscale interactome GCN and the PolypharmDB drug repurposing database ^36^, as described in **Fig. 1** *and in Materials and Methods*. Drugs assert their function by binding to proteins and regulating biological pathways. Therefore, we first constructed a multiscale interactome network to represent the known relation of 1661 drug molecules, 17,660 human proteins, 9,798 functional pathways and finally the 26 expressed proteins from SARS-CoV-2. This representation enables fine grained analysis on the probable drugs and protein targets for COVID-19, based on the assumption that the targets interacting with the viral protein nodes on the network are more likely to play a role in the potential treatment. Thus, we proposed a combined approach of node2vec and GCN to learn and generate the node embeddings in the multi-scale network. The embedding is optimized in an unsupervised manner to the objective that related nodes on the network should have higher embedding similarity (See *Materials and Methods*). All embeddings are then sorted by the proximity to the COVID-19 node. When first reviewing proximity distances between COVID-19 and human targets, very few had known small molecule modulators. To overcome the sparsity of drug-target interaction in the network, off-target predictions for relevant low-data targets were retrieved from the PolypharmDB database. PolypharmDB is a database of precompiled all-by-all Drug Target Interaction (DTI) predictions performed by the MatchMaker deep-learning engine, for 8535 human proteins with 10,244 clinically-tested small molecules (See *Materials and Methods*).

**Figure 1.**
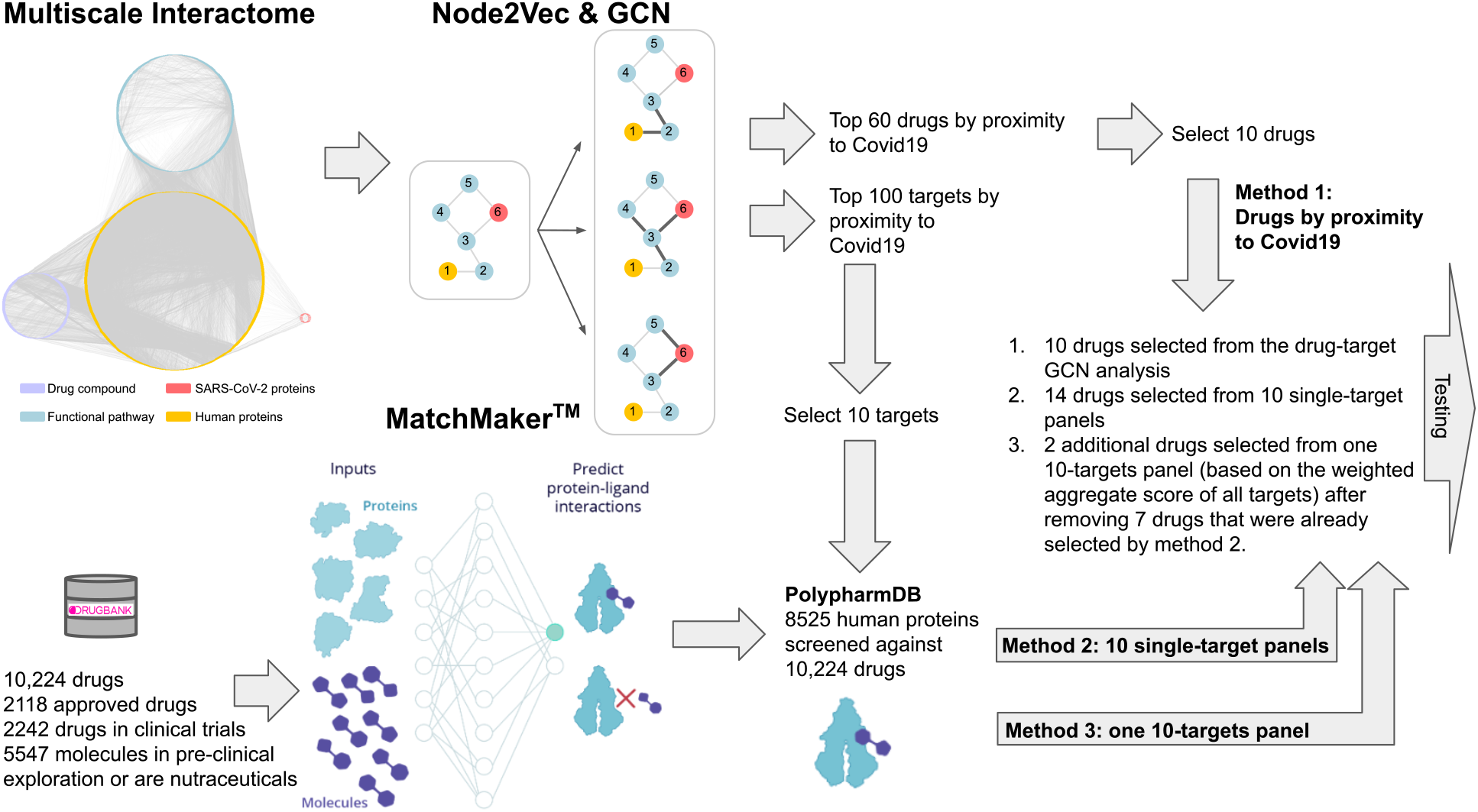
Three methods of selecting top drug candidates based on the GCN and MatchMakerTM prediction. Twenty-six drugs were selected for experimental testing by applying the predictions made by Node2Vec/GCN and MatchMakerTM models using three different methods. For *Method 1*, ten drugs were selected directly from top 60 drugs that were predicted to be most proximal to Covid19 based on the GCN alone. For *Methods 2* and *3*, first, ten human protein targets were selected from the list of top 100 proteins that are most proximal to COVID-19 based on the GCN prediction. Following that, for *Method 2*, 14 drug candidates were selected from top ten top ranking PolypharmDB (i.e., 10,224 drugs from DrugBank ^88^ screened against 8525 human proteins; see *Materials and Methods*) candidates for each protein (i.e. ten single-target panels resulting in 100 candidates in total). For *Method 3*, nine out of the top 25 ranking PolypharmDB candidates for the single ten-targets panel were selected as candidates, seven of which were already present in the list of candidates selected with *Method 2*. Please see Table 1 and Materials and Methods for further details.

For *Method 1*, the top 10 drug candidates were selected among the 60 drugs with the highest proximity scores to COVID-19, as determined by the GCN approach. For *Method 2*, 14 candidates were selected among single-target PolypharmDB hits for protein targets with high proximity scores to COVID-19 as determined by the GCN approach. For *Method 3*, the final two candidates were selected with a variation on *Method 2*, which prioritises compounds with multiple predicted interactions to GCN-identified SARS-CoV-2 targets. The compounds selected using all three methods are summarized in **Table 1**.

**Table 1.** Drugs that were selected for testing using three methods based on GCN analysis and MatchMaker™ predictions.*

| Method 1: Drug-target GCN analysis |  | Method 2: PolypharmDB query for 10 single GCN-target panels |  | Method 3: PolypharmDB query for one 10 GCN-targets panel |  |
| --- | --- | --- | --- | --- | --- |
| Drug | Target | Drug | Target | Drug | Panel of Targets |
| Bucladesine<br>Cinnarizine<br>Doxycycline<br>Eflornithine<br>Flucytosine<br>Glyburide<br>Nelarabine<br>Opicapone<br>Otilonium<br>Simvastatin | PRKACA<br>CACNA1C<br>PADI4<br>SLC25A21<br>DNMT1<br>ABCA1<br>POLA1<br>COMT<br>CACNA1C<br>ITGB2 | Bortezomib<br>Cefotiam<br>Dapagliflozin<br>Degarelix<br>Fosamprenavir<br>Gentamicin<br>Glipizide<br>Palbociclib<br>Polidocanol<br>Saquinavir<br>Streptozocin<br>Sugammadex<br>Telithromycin<br>Tofacitinib | IRAK4<br>TARS2<br>UGGT2<br>CD46<br>ADAM15<br>HEPACAM<br>ADAM15<br>IRAK4<br>MOGS<br>CD46<br>HEPACAM<br>MOGS<br>CD46<br>IRAK4 | Anidulafungin<br>Capmatinib<br>*Bortezomib*<br>*Dapagliflozin*<br>*Fosamprenavir*<br>*Glipizide*<br>*Palbociclib*<br>*Streptozocin*<br>*Tofacitinib* | UGGT2, SDF2,<br>NLRX1,<br>MOGS,<br>HEPACAM,<br>IRAK4,<br>ADAM15,<br>CD46, LILRA3,<br>and CHPF2 |

\*Top 26 drug candidates were selected based on GCN and MatchMaker predictions as described in Fig. 1 and in Materials and Methods. Cefotiam, under Method 2, was selected from the list of 10 top ranking candidates for TARS2, which was among the top 100 human proteins predicted to be associated with COVID-19 through target-only GCN analysis, but was not prioritized in the initial selection of top 10 targets described above. Drug candidates that were selected using Method 3 and were also predicted by Method 2 are shown in grey font and italics.

Since this approach generates numerous potential hits for further validation, we chose to use a system that could be widely used and scalable without restricted access to BSL3 laboratories, as required for monitoring of SARS-CoV-2 infection. This method may bias for identification of drug compounds with pan-human coronavirus antiviral activity, rather than compounds with selective antiviral activity towards SARS-CoV-2. Thus, to examine whether the 26 selected compounds exhibit antiviral activity, we established a cell culture-based immunofluorescence (IF) screening system using the 229E human alphacoronavirus, based on detection of the 229E S protein. This assay was designed to allow ~100% of control cells to express the S protein following 48 h of infection, allowing robust detection of antiviral activity as reduction of S protein abundance. We validated this IF assay by treating 229E-infected cells with the nucleoside analogue prodrug remdesivir ^27^ (**Fig. S1**), showing that this BSL2-based screening system allows for the safe and straightforward identification of novel antiviral compounds.

Using this assay, we identified four putative antivirals within the 26 predicted hits (4/26, ~15%) that reduced S protein abundance following 229E infection by at least 50% (**Fig. 2**) with no apparent cytotoxicity. Treatment with palbociclib and anidulafungin caused partial attenuation of 229E infection, while treatment with capmatinib or polidocanol resulted in nearly 100% inhibition of 229E infection. Based on our *in silico* framework, these four compounds were predicted to target several host proteins, each with potential novel and noncanonical roles in supporting coronavirus infection (**Table 1**). Interestingly, palbociclib, capmatinib and anidulafungin were predicted to target IRAK4, either as a primary target or by a polypharmacology panel. Another compound, bortezomib, was also predicted to target IRAK4, however cytotoxicity prevented further analysis of this compound.

**Figure 2.**
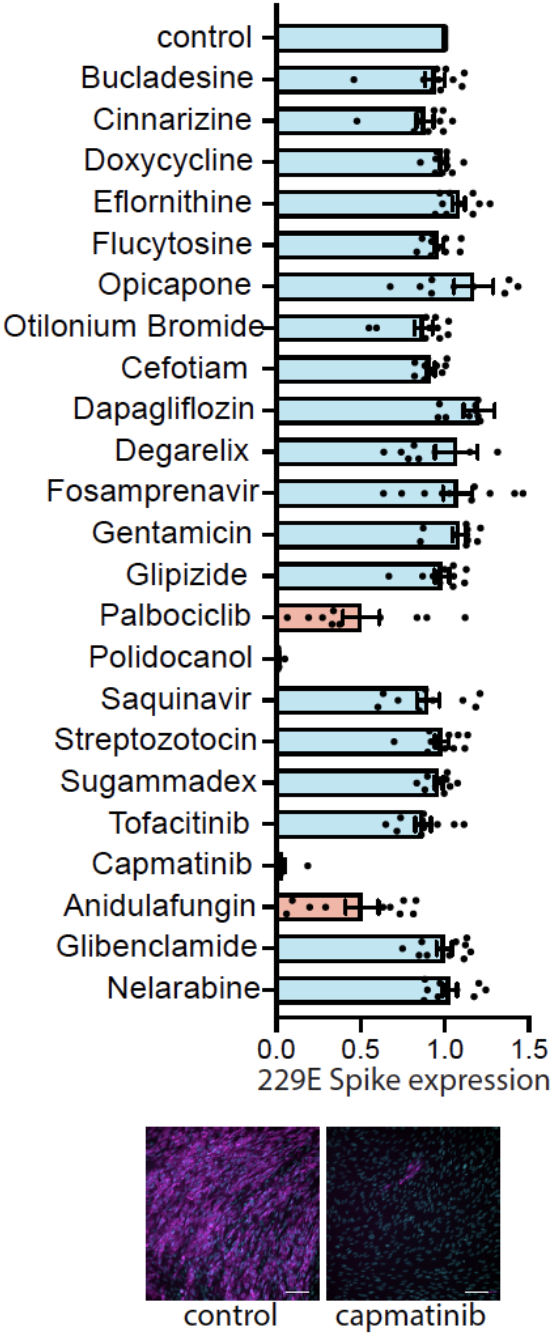
Screening of predicted compounds identifies capmatinib and other drugs as a host-targeted compounds with antiviral activity against human coronaviruses. Graph depicting mean ± SE and individual measurements of 229E Spike protein expression as measured by IF assay upon incubation with drugs as per **Table S4**. Results are expressed as mean 229E Spike expression relative to the DMSO vehicle (control) condition. (bottom) Representative images showing S protein expression (magenta) or DAPI (cyan) of the DMSO vehicle (control) or capmatinib (10 μM) treated conditions. Scale, 100 μm.

We focused further investigation into the role of capmatinib as a potential antiviral therapeutic against human coronavirus disease, given its robust impairment of 229E infection in the IF assay. Capmatinib is an orally-available inhibitor of the receptor tyrosine kinase (RTK) MET, and is used in the clinic for the treatment of MET-amplified non-small cell lung cancer ^37^; however, our analyses predict that capmatinib may have antiviral activity due to inhibition of targets other than MET.

To further characterize the antiviral activity of capmatinib on human coronaviruses, we developed several complementary cell-based assays to probe the different aspects of infection by several different human coronaviruses, including 229E, OC43, and NL63. Noteworthy, capmatinib treatment impaired 229E viral replication in a dose-dependent manner, with concentrations as low as 1.0 μM resulting in significant attenuation of 229E S protein abundance following infection (**Fig. 3A**). The ability of capmatinib to impair viral replication was further confirmed using a plaque-forming unit (PFU) assay in MRC-5 cells. 229E infection of control cells resulted in the formation of distinct, large circular plaques indicating cytopathic effect (CPE) (**Fig. 3B**). In contrast, capmatinib treatment resulted in reduced total number and size of individual plaques; viral quantification revealed that capmatinib treatment resulted in a > 50% decrease in viral production compared to control. Importantly, this method for viral quantification is based entirely on total plaque number and does not consider plaque size, thus, these results likely represent an underestimate of the antiviral effects of capmatinib, given the differences in plaque morphology between control and treatment groups (**Fig. 3B**).

**Figure 3.**
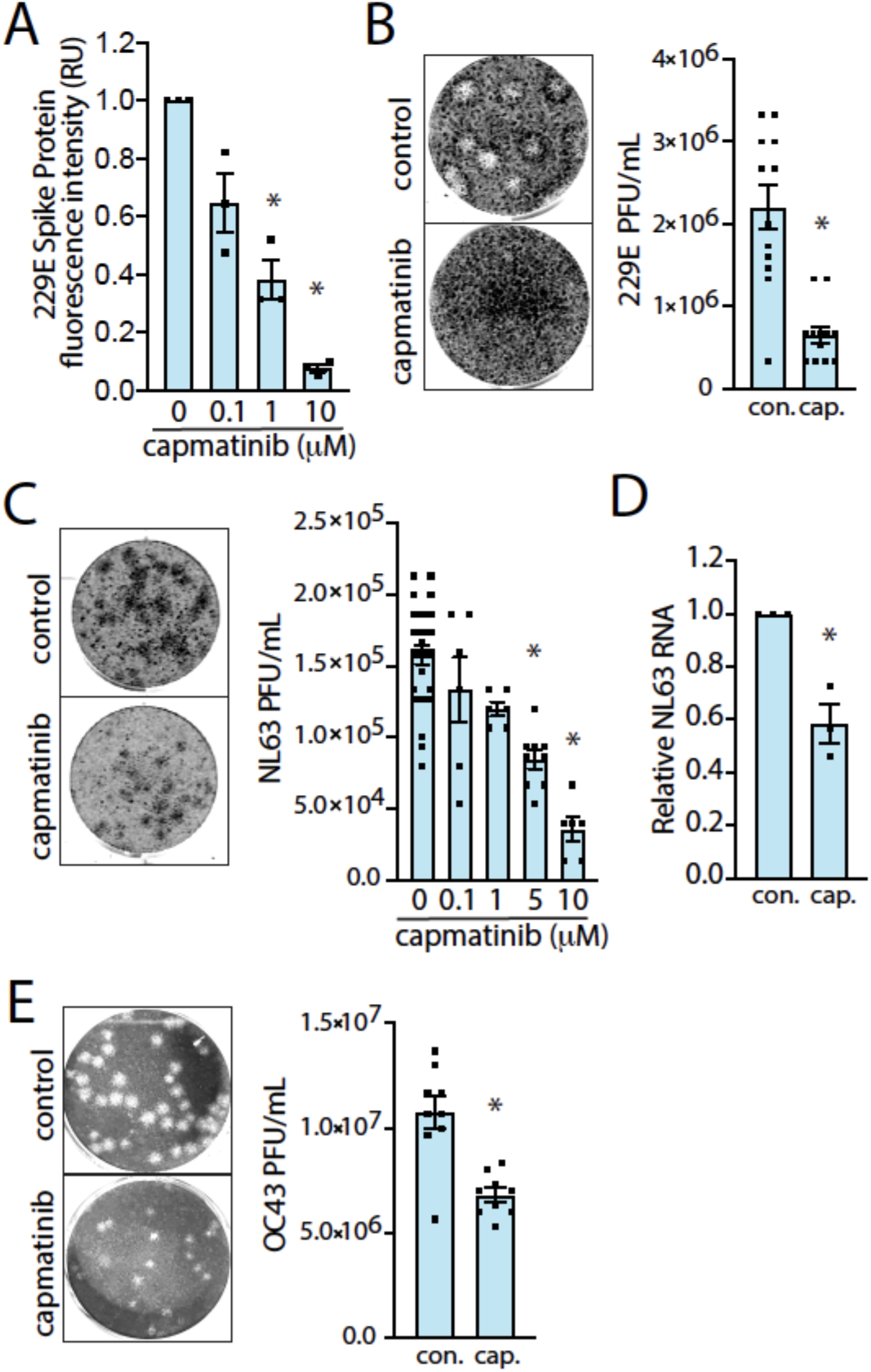
Capmatinib has a broad range of antiviral activity against human coronaviruses. A. Quantification of 229E Spike protein abundance in MRC-5 cells treated with increasing doses of capmatinib in the IF assay (48h infection), as mean ± SE (n = 3) expressed relative to DMSO (vehicle) control. *, p < 0.05 relative to control B. (left) Representative images of 229E plaques observed in MRC-5 cells treated with 10 μM capmatinib or DMSO (vehicle) for 6 days. (right) Quantification of viral titer from the DMSO (vehicle) control or capmatinib plaque assays, expressed as PFU/mL. *, p < 0.05 relative to control C. (left) Representative images of NL63 plaques observed in LLC-MK2 treated with 10 μM capmatinib or DMSO (vehicle) control for 5 days. (right) Quantification of NL63 PFU/mL in LLC-MK2 cells treated with the indicated doses of capmatinib. *, p < 0.05 relative to control D. Relative NL63 N protein RNA abundance 3 days post-infection in LLC-MK2 cells treated with 10 μM capmatinib relative to DMSO (vehicle) control. *, p < 0.05 relative to control, n=3 experimental replicates. E (left) Representative images of OC43 plaques observed in LLC-MK2 cells treated with 10 μM capmatinib or DMSO (vehicle) for 5 days. (right) Quantification of OC43 PFU/mL in LLC-MK2 cells treated with capmatinib or DMSO (vehicle) control. *, p < 0.05 relative to control.

To determine the breadth of the antiviral effects of capmatinib, we measured whether capmatinib also exhibited antiviral activity for other BSL2 human coronavirus infections, including OC43 and NL63. We adapted our MRC-5/229E plaque assay protocol for infection of LLC-MK2 cells with the alphacoronavirus NL63, which like SARS-CoV-2 depends on ACE2 for infection ^8^. Treatment of LLC-MK2 cells with increasing concentrations of capmatinib resulted in a dose-dependent decrease in NL63 PFU with minimal cytotoxicity observed at 5 days post-infection (dpi), with >70% reduction in PFU at 10 μM capmatinib (**Fig. 3C**). Consistent with results obtained with the PFU assay, qRT-PCR analysis showed that capmatinib attenuated NL63 N RNA abundance by approximately 50%, measured at 3 dpi (**Fig. 3D**). We also established a PFU assay based on the infection of LLC-MK2 cells with human betacoronavirus OC43. Treatment of OC43-infected LLC-MK2 cells with capmatinib resulted in a nearly 50% reduction in viral particles (**Fig. 3E**). As with our results obtained in the MRC-5/229E model, capmatinib treatment reduced both plaque number and the overall size of the plaques, indicating that our quantification of PFU/mL likely underestimated the actual antiviral effect of the drug. Taken together, our results from 3 different human coronaviruses and cell-based assays of coronavirus infection demonstrate that capmatinib has a broad range of antiviral activity against human coronaviruses.

We next explored the mechanism of the antiviral action of capmatinib. To determine whether the potent antiviral activity of capmatinib was due to its canonical role as a MET inhibitor, we compared the effects of capmatinib with another distinct MET inhibitor, AMG-337 that has similar *in vitro* IC_50_ ≤ 1 nM for MET as capmatinib ^38,39^. We first treated NL63-infected LLC-MK2 cells with capmatinib or an equimolar amount of AMG-337. As expected, capmatinib treatment greatly attenuated CPE as measured by PFU (**Fig. 4A**). In contrast, AMG-337 treatment, while similarly tolerated by the cells, did not result in an appreciable reduction in PFU. Similar results were observed in OC43-infected LLC-MK2 cells and 229E-infected MRC-5 cells; AMG-337 treatment had no effect on PFU in either case (**Fig. 4B** and **Fig. S2**). Capmatinib, but not AMG-337 was also effective at reducing 229E S protein abundance in the IF assay (**Fig. 4C**).

**Figure 4.**
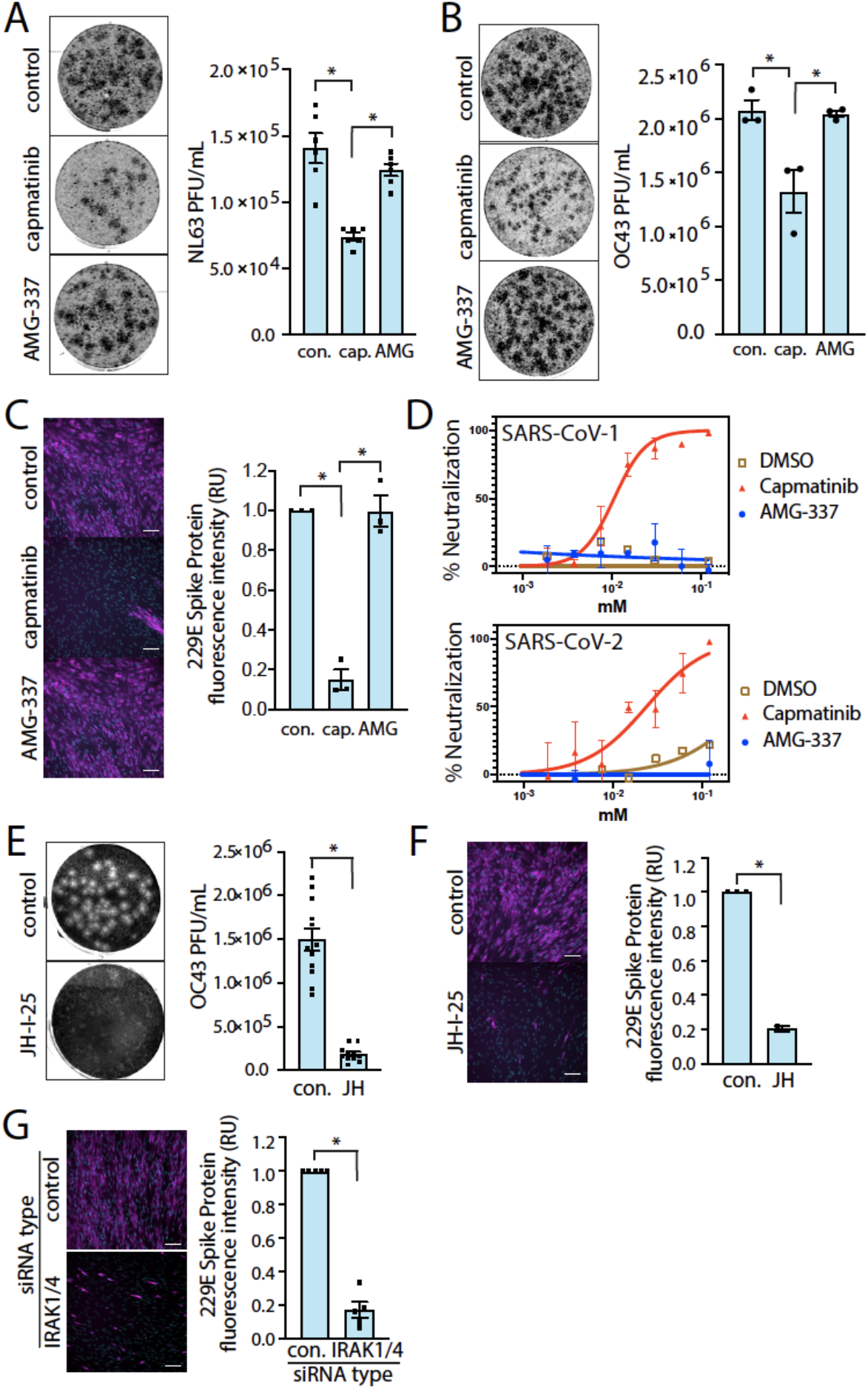
The antiviral activity of capmatinib is not attributed to its canonical role as an inhibitor of MET. A (left) Representative images of plaques in LLC-MK2 cells treated with 10 μM capmatinib, 10 μM AMG-337, or DMSO (vehicle) control and infected with NL63 for 5 days and (right) quantification of NL63 viral titer, shown as mean PFU ± SE (n = 3 with 3 technical replicates per experiment). B (left) Representative images of plaques in LLC-MK2 cells treated with DMSO (vehicle) control, 10 μM capmatinib, or 10 μM AMG-337, and infected with OC43 and (right) quantification of OC43 viral titer shown as mean PFU ± SE (n = 3 with 3 technical replicates per experiment). C (left) Representative images from IF assay of MRC-5 cells treated with 10 μM capmatinib or AMG-337 and infected with 229E. (right) Quantification of 229E S protein expression (20 images per condition, n=3). D. Representative neutralization curves from n=5 independent experiments showing the relative antiviral activity of capmatinib vs. AMG-337 in pseudovirus assays performed with the SARS-CoV-1 and SARS-CoV-2 Spike protein. E (left) Representative images of plaques in LLC-MK2 cells treated with 10 μM JH-I-25 and infected with OC43 for 5 days and (right) quantification of the OC43 viral titer shown as mean PFU ± SE (n=3 with 3 technical replicates per experiment). F (left) Representative images from MRC-5 cells treated with 10 μM JH-I-25 and infected with 229E for 2 days and (right) quantification of 229E Spike protein expression (10 images per condition, n=3). G (left) Representative images of MRC-5 cells transfected with IRAK1/4 siRNA or control and infected with 229E for 2 days. (right) Quantification of 229E Spike protein expression shown as mean ± SE, expressed relative to DMSO (vehicle) control (n=3). *, p < 0.05.

Spike-pseudotyped viral inhibition assays also demonstrate the ability of capmatinib to interfere certain aspects of viral infection. SARS-CoV-1 and SARS-CoV-2 pseudoviruses (PsV) require human ACE2 for cell entry ^40^. Incubation of HeLa cells stably expressing ACE2 (HeLa-ACE2 cells) with capmatinib showed a potent inhibition of infection of both SARS-CoV-1 or SARS-CoV-2 PsV with half-inhibitory concentrations (IC_50_) of 14.1 ± 0.2 and 26.0 ± 0.1 μM, respectively (**Fig. 4D**). In contrast, treatment with concentrations of AMG-337 up to the mM range had no appreciable effect on PsV neutralization (**Fig. 4D**). Both compounds displayed minimal cytotoxicity in HeLa cells (**Fig. S3**). Together, these results indicate that capmatinib exhibits human coronavirus antiviral activity by inhibiting a target other than MET. Moreover, the pseudotyped virus assay selectively monitors virus uptake and reporter gene delivery, suggesting that capmatinib inhibits early stages of virus entry.

We next sought to gain insight into the possible mechanism by which capmatinib exhibits human coronavirus antiviral activity. Based on MatchMaker, capmatinib was predicted to exhibit off-target interactions with IRAK4 and other protein targets (**Table 1**), which are part of pathways (e.g. interleukin-1 receptor) that have been broadly implicated in mediating SARS-CoV-2 infection and COVID-19 disease ^41,42^. In addition, our *in silico* analyses also predicted palbociclib and anidulafungin as binders of IRAK4, and as these drugs also demonstrated antiviral activity (**Fig. 2**), this suggests that IRAK4 may be an important novel drug target for human coronavirus infection and that capmatinib and other drugs may exhibit antiviral activity by novel action on IRAK4.

IRAK4 (Interleukin-1 receptor-associated kinase 4) is a serine/threonine kinase that forms part of the myddosome complex (consisting of MyD88 and other IRAK kinases) to transduce signals from Toll-like receptors (TLRs) or interleukin receptors ^43–45^. In canonical myddosome signal transduction, activation of membrane TLRs results in recruitment of the adaptor protein MyD88 and its associated IRAKs. MyD88-bound IRAK4 recruits and phosphorylates additional IRAK kinases, such as IRAK1 and IRAK2, which can then dissociate from the myddosome and trigger a signaling cascade involving subsequent TRAF6 and TAK-1 activation ^43–45^. These signaling events ultimately result in activation of the transcription factors NF-κB and MAPKs, which together contribute to the innate immune response to pathogens. Importantly, IRAK4 exhibits some functional redundancy with the related kinase IRAK1 ^46^, which led us to consider inhibition of IRAK1 and IRAK4 as an antiviral mechanism for human coronavirus infection.

We examined the effect of JH-I-25, a dual specific IRAK1 and IRAK4 inhibitor with no clinical relevance with IC_50_ values of 9.3 nM and 17.0 nM ^47^ respectively, on human coronavirus infection. Treatment of LLC-MK2 cells with JH-I-25 (10 μM) greatly reduced CPE in cells infected with OC43 (**Fig. 4E**). Similar results were obtained with the IF assay in MRC-5 cells infected with 229E (**Fig. 4F**). To verify that our results were specific to the combined inhibition of IRAK1 and IRAK4, we silenced both using siRNA gene silencing. In concordance with results obtained from the JH-I-25 experiments, combined silencing of IRAK1 and IRAK4 attenuated 229E S protein abundance (**Fig. 4G**). Consistent with a role for IRAK1/4 in coronavirus infection, inhibition of p38 MAPK, known to be activated downstream of IRAK1/4 ^43–45^, also impaired coronavirus infection (**Fig. S4**). Taken together, these data demonstrate that the effects of combined IRAK1/4 inhibition recapitulate the effects of capmatinib, as predicted by our *in silico* analyses.

## Discussion

### Computational analysis and antiviral protein target predictions

*In silico* drug repurposing approaches have the innate advantage of accessing very large collections of information to uncover indirect associations that may have been otherwise overlooked. In the case of target-identification tasks, broadening the search of plausible targets invariably introduces a new challenge, as the overwhelming majority of human proteins have not been explicitly targeted by existing small molecule drugs. In this study, we addressed these challenges simultaneously by pairing two ML approaches designed for target-identification and drug-target interaction predictions, demonstrating the viability of an *in silico* first repurposing workflow, coupled with robust bioactivity assays

We integrated drug-target interactions, proteins and functional pathways in a multiscale interactome network. Based on the assumption that drugs take effect by binding to proteins and regulating pathways, the multiscale interactome traces the biological processes of available treatments via the interactions across proteins, functional pathways, drugs and the target disease, COVID-19. To capture this process, our approach used biased random walks and Graph Convolutional Network (GCN) to model the correlations of nodes of multiple types and build embeddings for each drug, protein and pathway. The embeddings were optimized in an unsupervised manner to encode the direct and indirect relations in the multiscale interactome. Then by ranking the proximity between the embeddings of candidate proteins to the target disease, the GCN model determined the potential efficacy of drugs or protein targets for this disease.

### Matchmaker, PolypharmDB, and drug repurposing

The multiscale interactome GCN unveiled multiple plausible viral-host targets for repurposing leads. Rather than subjecting a single protein to a target-centric computational screen, the subsequent screening staged maximized diversity in both targets and their putative binders. Repurposing targets prioritized by the GCN were cross-referenced to PolypharmDB, a precompiled database of off-target interaction predictions between 8525 human proteins and 10,244 small molecule compounds with prior clinical evaluation. This approach contrasts sharply with disease-centric or target-centric approaches, which make up the overwhelming majority (~90%) of drug repurposing studies ^48^. PolypharmDB combines drug-centric and target-centric repurposing strategies by identifying novel off-target interactions across the human proteome with known structures *in silico*. Evidence that capmatinib’s antiviral activity is driven by interactions other than its known target MET is provided by the lack of antiviral activity of AMG-337, another potent inhibitor of MET RTK (**Fig. 4A-D**). AMG-337s is chemically distinct from capmatinib, which likely drives its differential polypharmacology.

Criteria were set to maximize the diversity of compound and target selection for downstream evaluation. The recognition that polypharmacology may play a role in effective therapeutics motivated the evaluation of multiple targeted agents, leading to the nomination of capmatinib and other compounds for testing. Underappreciated polypharmacology may play a larger role in the activity of small molecule drugs; a study investigating oncology medications found for several drugs investigated that on-target interactions were inconsequential for drug activity, with off-target effects driving efficacy ^49^. Alternatively, the use of multiple targets as separate or as combined objectives may have contributed to the success of this investigation simply by providing more possibilities for favorable outcomes. Nonetheless, there may be broader opportunities within a drug-centric approach for drug repurposing, where validated therapeutic targets can be linked to small molecule drugs through approaches like PolypharmDB.

### Capmatinib as an antiviral drug for COVID-19 and other human coronavirus diseases

Capmatinib was developed as a MET RTK inhibitor for the treatment of various MET-amplified tumors ^50^. The MET proto-oncogene encodes an RTK that serves as the receptor for hepatocyte growth factor (HGF). A number of small-molecule MET inhibitors have recently been developed and are currently undergoing clinical trials to determine their efficacy in reducing cancer morbidity and mortality ^51,52^. The availability of data on the safety of MET inhibitors in the clinic, and in particular capmatinib, provides support for this strategy of drug repurposing via polypharmacology.

We observed that capmatinib exhibited antiviral activity in the low micromolar range. Indeed, using the SARS-CoV-2 and SARS-CoV-1 pseudotype virus infection assay, we determined that the IC_50_ for capmatinib for neutralization of infection was 14.1 ± 0.2 and 26.0 ± 0.1 μM, which is in the range reported for action of other drugs targeting SARS-CoV-2 infection when tested in Vero and Calu-3 cells in culture, including remdesivir ^53^. Hence, capmatinib should be considered as a candidate for therapeutic testing in further preclinical and clinical trials for the treatment of human coronavirus diseases, including COVID-19. In addition, while we have not tested the action of capmatinib against SARS-CoV-2 variants that have emerged in 2021 ^6,7^, the broad antiviral activity that capmatinib exhibits against 5 genetically different human coronaviruses (229E, OC43 and NL63 live virus infection and SARS-CoV-1 and SARS-CoV-2 pseudotyped virus assays) suggests that capmatinib may hold promise in broadly treating SARS-CoV-2 variants of concern, or other variants that may arise from further antigenic drift as well as other emerging viruses. To our knowledge, this is the first demonstration of antiviral activity of capmatinib in cell-based coronavirus infection assays. While this manuscript was in preparation, other computational approaches predicted binding of capmatinib to the SARS-CoV-2 S protein, viral proteases, RNA-dependent RNA polymerase and/or viral endoribonuclease ^54–56^. While our analysis indicates that capmatinib may act by targeting IRAK signaling, and we find a novel requirement for IRAK1/4 in supporting human coronavirus infection, the mechanism of antiviral action of capmatinib warrants further investigation in future studies.

### Novel role for IRAK signaling in supporting human coronavirus infection

Myddosome signaling is considered an essential component of the innate immune response to bacterial PAMPs, and loss of function mutations in IRAK4 or MyD88 results in a primary immunodeficiency syndrome associated with a dramatic increase in susceptibility to specific pyogenic bacterial infections ^57–60^. In contrast, MyD88 and IRAK4 may have distinct roles in response to other pathogens, as their perturbation often does not impact susceptibility to some viral, fungal, protozoal, or other infectious agents ^57–61^. Notably, MyD88 perturbation contributes to coronavirus infection and symptom severity in animal models ^62^. However, whether and how human coronaviruses may engage TLR and IRAK signals during infection remains poorly understood. While IRAK1 and 4 are broadly expressed, in single-cell datasets deposited in the covid19atlas, IRAK1 and IRAK4 appear highly expressed in Secretory3 cells in the bronchial epithelial dataset ^63^. Interestingly, Secretory3 cells have the highest expression of ACE2 as compared to other cell types in the primary human bronchial epithelial cell dataset, consistent with a role for IRAK1/4 in supporting coronavirus infection.

Our results from two different human coronaviruses and two different host cell lines suggest that human coronavirus infection requires IRAK1 and/or IRAK4. We focused our studies on concomitant perturbation of IRAK1 and IRAK4, given the redundancy that has been reported between these kinases in some contexts ^64^. Consistent with our results, another study performed an analysis of phosphorylated sequences within host cells and predicted activation of IRAK4 within 15 min of infection of Vero cells with SARS-CoV-2 ^65^. Together with the results presented here, this suggests that IRAK1/4 may contribute a non-canonical function to support human coronavirus infection. Recent work has also established that modulation of IRAK4 dependent immune responses is crucial for mounting an appropriate immune response during SARS-CoV-2 infection, supporting this observation ^66^.

As such, therapeutic modulation of IRAK signaling, and perhaps also that of TLRs and MyD88, may be a useful strategy for treatment of patients with COVID-19. In fact, a phase II clinical trial is ongoing to probe the use of the IRAK4 selective inhibitor PF-06650833 to treat COVID-19 patients with acute respiratory distress syndrome ^67^. Supporting the role of this signaling pathway in the progression of the disease, obese individuals have increased TLR/MyD88 signaling that may predispose them to severe COVID-19 symptoms ^68^.

In this study, we use multiple methods involving a multiscale interactome GCN and the PolypharmDB drug repurposing database to identify new drug targets and drug repurposing candidates for the treatment of human coronavirus disease. We also provide evidence that several drug molecules predicted by this method have previously unknown antiviral activity against human coronavirus infection, in particular capmatinib. Further, we identify IRAK1/4 as new and unexpected coronavirus drug targets, required for coronavirus infection. This indicates that the methods described herein are a novel and powerful approach for the rapid identification of new therapeutic strategies to identify antiviral drugs and could also be applied more widely for novel therapeutic intervention to other classes of disease.

## Materials and Methods

### Materials

#### Viruses

229E and OC43 coronaviruses were obtained from the American Type Culture Collection (ATCC) (ATCC® VR-740™ and ATCC® VR-1558™). NL63 coronavirus was kindly provided by Dr. Scott Gray-Owen (University of Toronto). Original viral stocks were stored at −80°C until use and all subsequent viral stocks were produced from the original parental stocks.

#### Cell lines

MRC-5 (lung fibroblast) cells was obtained from ATCC (ATCC® CCL-171™). CoronaGrow LLC-MK2 (kidney epithelial) cells, a subclonal line of parental LLC-MK2 cells were obtained from VectorBuilder Inc. (Chicago, IL, Cat. No. CL0004). HEK293T cells expressing the full-length SARS-CoV-2 spike (BEI NR52310) were obtained from BEI Resources (Manassas, VA). HEK293T cells expressing the SARS-CoV-2 spike were kindly provided by S. Pöhlmann (Leibniz Institute for Primate Research, Göttingen, Germany). HeLa-ACE2 cells were kindly provided by D.R. Burton (The Scripps Research Institute).

### Multiscale Interactome

A base multiscale interactome network combining drug-protein interactions (8,568), human protein-protein interactions (387,626) and protein-pathway interactions (22,545) was retrieved from Ruiz et al. ^32^. The base network aggregates data from multiple primary sources ^69–79^. An additional 332 experimentally-derived, viral-host protein-protein interactions were added to adapt the network for SARS-CoV2 repurposing ^23^. Then for a given viral protein that interacts with human proteins, we investigated the pathways in which these human proteins are involved. We added direct connections between viral proteins and the retrieved human functional pathways. In our experiments, adding direct connections between viral proteins and human pathways shows significant improvement in performance. Lastly, COVID-19 was introduced as a final entity to the multiscale interactome network and linked to all SARS-CoV-2 proteins. The multiscale interactome was represented as a graph *G* = (*V, E*), where individual proteins, pathways and drugs form vertices (*V*) and interactions form the edges (*E*).

The goal of our approach is to learn meaningful embeddings of the nodes in the multiscale interactome network so that we can predict drug candidates or protein targets using these embeddings. To propose a proper embedding function, naively, we can introduce the bias of graph homophily, i.e. drugs/targets that connect to the viral protein via the shortest paths are the most likely to be effective. This is reflective of the assumption that drug effects propagate along the biological network to treat diseases. However, non-homophily relations can be equally important - for example, a protein can be a promising target if it is involved in several important related pathways, although it may not be directly connected to viral proteins. To address this challenge, we use graph convolutional networks to learn this hierarchy of complex relations from the interactome data.

#### Graph Convolutional Network

To prepare the Graph Convolutional Network (GCN), initial node embeddings were generated using Node2Vec ^80^. The return parameter *p* and “in-out” parameter *q*, in Node2Vec were set to 0.25 in order to balance global and local views of the random walk process ^81^, which helped capture the aforementioned homophily and non-homophily relations. The embedded dimensions size *D* was set to 64 and node2vec was applied to the multiscale interactome graph to convergence. The Node2vec output was an embedded matrix *H_n_* ∈ ℝ^*K,D*^, where *K* = |*V*| is the number of nodes, and each row in *H_n_* is the learned representation for the corresponding node.

These embeddings were used as the initial input feature layer, *H*^(0)^, in our GCN model. The model consists of stacked multiple Graph Convolution and each one of them is defined by equation 1 ^33^. In this model, *σ* is the activation function ReLU, 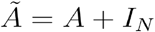 is a transformation of the multiscale interactome adjacency matrix (*A*) with the additional self-associating nodes (identity matrix *I_N_*), *H*^(*l*)^ and *W*^(*l*)^ representing layer-specific feature vectors and trainable weights, while 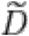 is a diagonal degree matrix defined by 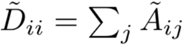,

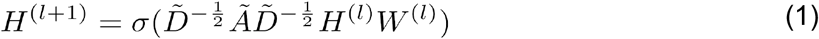

The adjacency matrix *A* ∈ ℝ^*K,K*^ in equation 1 is generated by the edges *E* in the multiscale interactome. Each element in *A* is either the connection weight or zero if there is no connection. The output features of the last layer 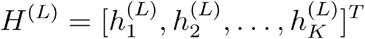 are then normalized to produce the final embedding for each node.

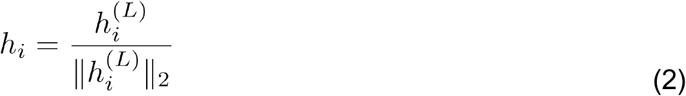

To train the GCN model, we first compute cosine similarity between the embeddings of all nodes, 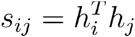 and use a diffusion loss inspired by Liu et al.^82^ to train the model parameters with gradient backpropagation,

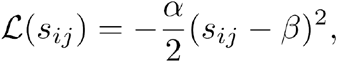

where *α* > 0, and *β* is a predefined threshold and set to 0.25 in this work.

The idea of the loss function is to cluster embeddings of relevant nodes in the multiscale network and separate those irrelevant embeddings conversely. This effect can be observed from the derivative of 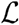:

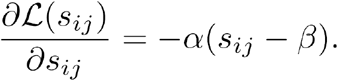

Therefore similarity values larger than threshold *β* are encouraged and embeddings of relevant nodes will cluster. Conversely, the loss function also diverges the embeddings of irrelevant nodes in the network even further away. Finally, we train the model for fixed 20 epochs and obtain the updated embeddings *h_i_* for each node. The final embeddings were then saved for further computation.

Lastly, the proximity values (i.e. cosine similarities) from COVID-19 to each drug and each human protein present in the multiscale interactome were calculated and ranked to identify direct repurposing candidates and plausible targets (see section “Compound Selection”).

### Drug-target Interaction Predictions

To identify small molecule drug candidates for host-based targets proposed by the GCN with few or no known binders, we looked up Drug Target Interaction (DTI) predictions in PolypharmDB ^36^. PolypharmDB is a drug-repurposing database of pre-computed drug-target interaction predictions, evaluated by using the 2020Q2 beta release of MatchMakerTM ^83^. Matchmaker is a deep learning model that predicts interaction of a small molecule drug/binding site pair using paired structural features of the drug with the 3D structural features of the protein binding sites. MatchMaker models with positive training examples complexes obtained by threading drug-target interaction (DTI) data ^84^ onto 3D structures of protein-ligand binding sites obtained from the Protein DataBank ^85^ and SwissModel ^86^, on the basis of chemical similarity ^83^. Negative training examples are obtained by random shuffling of positive DTI pairs, and models are trained on progressive thresholds of increasing stringency in accordance to the confidence in the source DTI and in the selection of representative 3D structures ^87^.

The compound screening library consists of 10,244 small molecule drugs, retrieved from DrugBank ^88^ on February 19th, 2020. The library excluded compounds with fewer than four carbon atoms, or whose SMILES chemical structure was unable to be parsed by RD-KIT (Release_2018_09_1) RDKit: Open-source cheminformatics; http://www.rdkit.org) ^89^. The screening set includes 2118 approved drugs, 2242 drugs in clinical trials and 5547 molecules in preclinical development or nutraceuticals.

The protein screening library represents 29,290 pockets from 8525 human proteins obtained by the PDB ^85^ (retrieved January 2018) and SwissModel ^86^ database (Retrieved June 2018). Specifically, pockets correspond to binding sites of drug-like molecules observed as cocrystalized ligands in the PDB source files, or superimposed ligands from template structures from SwissModel source files. Drug-target interaction pairs were evaluated using all combinations of small molecule drugs and available binding site structures. Individual human proteins were ranked on the basis of their top-scoring pockets. Additional details related to the construction of PolypharmDB and screening libraries are available from Redka et al. 2020 ^36^.

### Compound Selection

Small molecule repurposing candidates were selected from three methods combining the GCN network and PolypharmDB (**Fig. 1**) approaches. For each method, a systematic selection criterion was applied to maximize the diversity of assayed compounds, relevance of the hit to the infectivity-based assay, availability and repurposing appropriateness. Selected drugs for all three methods are provided in **Table 1**.

For *Method 1*, compounds were selected among drug candidates suggested directly by the GCN, on the basis of their network distance to COVID-19. Since the GCN input multiscale interactome contained a broad variety of drugs, we limited the selection to FDA-approved, small molecule drugs, selecting only the top hit for each family of structurally- or functionally-related compounds. All 10 drug candidates selected by this method were among the raw to 60 hits prior to applying the filter process.

*Methods 2* and *3* combined the GCN for viral-host target selection and PolypharmDB to predict small molecule binders for the proposed targets. Target selection was performed by ranking human proteins in accordance with their GCN proximity scores to COVID-19. The top 100 human proteins of 17,660 represented in the network were considered in the selection process. Proteins with a known or likely association to the adaptive immune response were excluded, as any tentative effect would not be measurable in a cell infectivity assay. Target selection was also restricted to one target per homologous family to diversify the screen. The final selected targets were as follows: UGGT2, SDF2, NLRX1, MOGS, HEPACAM, IRAK4, ADAM15, CD46, LILRA3, and CHPF2. *Methods 2* and *3* diverge in their compound selection process from PolypharmDB. For *Method 2*, the top ten repurposing hits for each of the ten targets were subject to an expert selection process. A total of 14 compounds from these lists were selected to maximize functional and structural diversity, as well as availability, prediction scores, and FDA approval status. *Method 3* ranked all PolyphamDB compounds according to a weighted aggregate score combining all ten GCN targets. The weighted aggregate score represents a probability of interacting with any or multiple GCN-selected targets. The top 25 ranked compounds were subject to the same expert selection process as *Method 2*, leaving 9 possible repurposing candidates. However, seven of these nine had already been selected by *Method 2*, leaving only two repurposing candidates in this category.

### Cell Culture and Human coronavirus propagation

For propagation of cell lines for use in coronavirus infection experiments, cells were maintained in a standard tissue culture incubator maintained at 37°C with 5% CO_2_. For infection of cells and propagation of coronaviruses, cells were maintained in a standard cell culture incubator maintained at 33°C with 5% CO_2_. This temperature was determined to support optimal propagation of human coronaviruses and yielded higher viral titers than preparations of the virus grown at 37°C.

MRC-5 and LLC-MK2 cell lines were maintained in *growth medium*, consisting of Minimum Essential Medium Eagle (MEM), with Earle’s salts, L-glutamine and sodium bicarbonate (Sigma Aldrich Cat. No. M4655) and supplemented with 10% heat-inactivated fetal bovine serum (FBS, ThermoFisher Scientific Cat. No. 10082147) and 1X penicillin-streptomycin (P/S, ThermoFisher Scientific Cat. No. 15070063). For all experiments involving coronavirus infection, cells were maintained in *infection medium*, which is the same as *growth medium*, except the concentration of heat-inactivated FBS was 2%. For plaque assays, cells monolayers were overlaid with *plaque media*, consisting of 2X MEM Temin’s modification, no phenol red (ThermoFisher Scientific Cat. No. 11-935-046), 2% heat-inactivated FBS, 1X P/S, and mixed 1:1 with pre-warmed 0.6% agarose (ThermoFisher Scientific Cat. No. 16500-500). Alternatively, see **Table S1** for specific formulations of culture media.

Propagation of human coronaviruses was performed based on suppliers’ instructions. Briefly, MRC-5 or LLC-MK2 cells were grown to 90% confluence in a T75 tissue culture flask in standard growth medium. Cell monolayers were washed 2X with infection medium prior to infection. Cell monolayers were infected with 229E (MRC-5 cells), OC43 (LLC-MK2), or NL63 (LLC-MK2) at a MOI of 0.01 in a total volume of 4 mL infection medium for 2 hours adsorption at 33°C with 5% CO_2_. Following viral adsorption, unbound virus was aspirated, and cell monolayers were washed 2X with infection medium, and then 12 mL fresh infection medium was added. Flasks were then placed back in the 33°C incubator for a period of 2-4 days depending on the coronavirus strain to achieve maximal viral titer. To harvest the virus, supernatant was collected in a 15 mL conical tube and centrifuged at 1000 x g for 10 min to pellet cell debris. Viral stocks were stored as single use aliquots at −80°C. Viral concentration of new preparations of viral stocks were measured by PFU assay.

### Immunofluorescence detection of 229E infection

Briefly, MRC-5 human lung fibroblast cells (seeded on glass coverslips) were infected with 229E at a MOI of 0.01 in the presence of drug (1-10 μM) or vehicle control for 1 h at 33°C and 5% CO_2_. Following incubation, excess unbound virus was removed, and cells were incubated in fresh infection medium with drugs for an additional 48 h. This time point was chosen because it was initially determined to result in infection of ~100% of control cells and thus provided a baseline for examination of antiviral activity. This time point was also chosen because the infected cells remained in a virtually intact monolayer with minimal CPE - extensive CPE could lead to sampling errors and could be confounded with potential cytotoxic effects of the drugs. Drugs that caused signs of cytotoxicity (identified by DAPI staining) were excluded from further analysis as they confounded interpretation of potential antiviral effects in the IF assay.

For the IF protocol, cells infected with coronaviruses were washed 2X with PBS (with Mg^2+^ and Ca^2+)^ and then immediately fixed for 1 h with 4% paraformaldehyde. This was followed by 15 min treatments with: 0.15% glycine, 0.1% triton X-100, and 3% bovine serum albumin with PBS washes in between each step. 229E S protein was detected with treating the cells with 100 μL (1:50 dilution) of the mouse anti-229E S protein antibody 9.8E12 ^90^ by the inverted drop technique for 1 h at room temperature. Following primary antibody incubation, cells were washed and treated with AlexaFluor488-conjugated anti-mouse secondary antibody (1:1000 dilution) (Cedarlane Labs Cat. No. 115-545-003) and DAPI (1 μg/mL) for 1 hour at room temperature. Following another wash, coverslips were mounted on glass slides with DAKO mounting media (Agilent Technologies Cat. No. S3023) and then incubated at room temperature overnight to solidify. Slides were visualized on an inverted microscope by widefield epifluorescence (Leica DMi8 microscope, Andor Zyla 4.2-megapixel camera, run by Quorum WaveFX by Metamorph software). For each IF experiment, a total of 10-20 randomly chosen fields were selected in the DAPI channel for acquisition of the 229E S protein with a 10x objective lens. Images were quantified by measuring the total fluorescence signal using ImageJ software (National Institutes of Health, Bethesda, MD) ^91^.

#### Human coronavirus plaque assays

All coronavirus plaque assay protocols were adapted from previously described methods for measuring viral concentration using PFU assays ^92–94^. Briefly, cells were grown to confluency on 6-well tissue culture plates and then infected with serially diluted virus in a volume of 300 μL for 1 h adsorption at 33°C with 5% CO_2_, with gentle agitation every 15 min. Following adsorption, unbound virus was removed, and cells were washed 2x with infection media and then overlaid with plaque media. In experiments involving the testing of potential antiviral compounds, drugs were added to the infection media during adsorption and to the plaque media. Following an incubation period of several days to establish CPE (see below for virus-specific information), cells were fixed with 10% neutral buffered formalin overnight, followed by removal of the agarose plug and counterstaining with 1% crystal violet solution to visualize the plaques. For all plaque assays, each condition was performed in technical triplicates.

##### 229E

Confluent monolayers of MRC-5 cells were grown on 6-well tissue culture plates and infected with serially diluted 229E virus. After washing away unbound virus, the cells were overlaid with a semi-solid agarose medium to restrict the spread of virus to adjacent cells. After 5-7 days incubation, cells were fixed and stained to quantify the number of plaques in each well.

##### NL63

NL63 viruses were propagated in the monkey kidney epithelial cell line LLC-MK2, which has been reported to support NL63 production ^95^. Confluent monolayers of LLC-MK2 cells were grown on 6-well plates and infected with serially diluted NL63 virus. After washing away unbound virus, the cells were overlaid with a semi-solid agarose medium to restrict the spread of virus to adjacent cells. After 4 days incubation, cells were fixed and stained to quantify the number of plaques in each well.

##### OC43

We determined that OC43 viruses could be readily propagated in LLC-MK2 cells, similar to NL63 viruses. Confluent monolayers of LLC-MK2 cells were grown on 6-well plates and infected with serially diluted OC43 virus. After washing away unbound virus, the cells were overlaid with a semi-solid agarose medium to restrict the spread of virus to adjacent cells. After 4 days incubation, cells were fixed and stained to quantify the number of plaques in each well.

### qRT-PCR detection of NL63

LLC-MK2 cells seeded on 6-well tissue culture plates were infected with NL63 at a MOI of 0.01 for a 1 h adsorption period in the presence of 10 μM capmatinib in a volume of 300 μL. Following viral adsorption, cells were washed 2x with infection media and then 500 μL fresh media/drug was added to the cells for a 3-day incubation period. To extract total NL63 RNA, 1 mL TRIzol™ Reagent (ThermoFisher Scientific) was added directly to the samples (cells + supernatant) and cells were lysed by scraping. Total RNA was purified using the Direct-zol™ RNA Miniprep Kits (Zymo Research, Irvine, CA) according to manufacturer’s instructions. Reverse transcription and qPCR were performed in a one-step reaction using Luna® Universal One-Step RT-qPCR (NEB, Ipswich MA) according to manufacturer’s instructions. RT-qPCR reactions were performed on a CFX96 thermal cycler (Bio-Rad, Mississauga, ON). Experiments were performed with at least 2 technical replicates to monitor variation between wells, no template/no RT controls, and melt curves. Reactions were performed at a final volume of 20 μL with 20 ng input RNA and a primer concentration of 500 nM. All results were normalized to GAPDH RNA levels and to the infected, non-drug treated condition. Relative change in NL63 RNA was calculated using the 2^-ΔΔCt^ method ^96^. A minimum of 3 experimental replicates were used to assess NL63 Nucleocapsid RNA using previous established primers ^97^. For a complete list of qPCR primers see **Table S2**. For thermal cycler conditions see **Table S3**.

### siRNA transfection

siRNA transfection was performed by two individual transfections at 72h and 48h prior to infection or other experimental manipulation, respectively. Briefly, MRC-5 cells were grown to 40% confluency before the first round of transfection. On the day of transfection, cells were placed in fresh growth medium. A transfection master mix was made in Opti-MEM (ThermoFisher Cat. No. 31985070) by diluting stock siRNA solutions to a final concentration of 50 nM and 6.3 μL Lipofectamine RNAiMAX transfection reagent (ThermoFisher Cat. No. 13778030) per well of a 6-well tissue culture dish. Cells were transfected for 4 h before switching to fresh growth medium. For a list of siRNA sequences see **Table S2**.

### Pseudotyped virus particle assay

SARS-CoV-2 pseudotyped viruses (PsV) were prepared using an HIV-based lentiviral system as previously described ^11^ with few modifications. Briefly, PsVs were produced by transfection of human kidney HEK293T cells with the full-length SARS-CoV-2 spike (BEI NR52310) or SARS-CoV-1 spike (kindly provided by S. Pöhlmann, Leibniz Institute for Primate Research, Göttingen, Germany). Cells were co-transfected with a lentiviral backbone encoding the luciferase reporter gene (BEI NR52516), a plasmid expressing the Spike (BEI NR52310) and plasmids encoding the HIV structural and regulatory proteins Tat (BEI NR52518), Gag-pol (BEI NR52517) and Rev (BEI NR52519). After 24 h at 37°C, 5 mM sodium butyrate was added to the media and the cells were incubated for an additional 24-30 h at 30°C. Next, the PsV particles were harvested, passed through 0.45 μm pore sterile filters and finally concentrated using a 100K Amicon (Merck Millipore Amicon-Ultra 2.0 Centrifugal Filter Units).

Neutralization was determined in a single-cycle neutralization assay using HeLa-ACE2 cells (kindly provided by D.R. Burton; The Scripps Research Institute). To that end, 50 μL of 2-fold serial dilutions of the small molecules were incubated with 10,000 cells/well seeded the day before (100 μL/well) for 1h at 37°C. After 1 h incubation, 50 μL of PsVs was added to each well and incubated for 48h-60h in the presence of 10 μg/mL of polybrene (Sigma Aldrich, TR-1003-G). Infection levels were inferred from the amount of luminescence in relative light units (RLUs) after adding 50 μL Britelite plus reagent (PerkinElmer) to 50 μL of media containing cell (i.e. after removing 130 μL/well to account for evaporation). After 2 min incubation, the volume was transferred to a 96-well white plate (Sigma-Aldrich) and the luciferase intensity was read using a Synergy Neo2 Multi-Mode Assay Microplate Reader (Biotek Instruments). Two to three biological replicates with two technical replicates each were performed. Culture media was prepared by supplementing DMEM media with 2% inactivated FBS and 50 μg/ml of gentamicin. IC_50_ values were calculated using Prism.

In order to confirm that the reduced infection was not related to cell toxicity, HeLa-ACE2 cell viability upon incubation with serial dilutions of the small molecules was assessed. 10,000 cells/ well of pre-seeded HeLa-ACE2 cells were co-cultured with 2-fold serial dilutions of the small molecules at 37 °C for 48h-60h under the same conditions as in the neutralization assay. Cell viability was monitored by adding 50 μL of CellTiter-Glo 2.0 reagent (Promega) to 200 μL of media containing cells. After 10 min incubation, 100 μL volume was transferred to a 96-well black plate (Sigma-Aldrich) to measure luminescence in relative light units (RLUs) using a Synergy Neo2 Multi-Mode Assay Microplate Reader (Biotek Instruments).

### Drug library

Unless indicated otherwise, all compounds were purchased from MedChemExpress (Monmouth Junction, NJ) and were reconstituted according to manufacturer’s specifications. A complete list of compounds is provided in the **Table S4**. All compounds were reconstituted to stock concentrations of 1-10 mM and frozen in individual aliquots at −80 °C until use.

### Statistical Analysis

All statistical analysis for biological experiments were performed with GraphPad Prism 9 software using student t-tests when comparing two conditions (**Figs 3B, 3D, 3E, 4B, 4E, 4F, 4G, S2A**) or one-way ANOVA with Tukey post-hoc test when comparing multiple conditions (**Figs. 3A, 3C, 4A, 4C, S4**).

## Acknowledgements

We gratefully acknowledge funding that supported this research support from the Ryerson University Faculty of Science (CNA), as well as funding support in the form of a CIFAR Catalyst Grant (JPJ and CNA), an NSERC Alliance Grant (CNA) and the Ryerson COVID-19 SRC Response Fund award (CNA). BW is partly supported by CIFAR AI Chairs Program. This work was also supported by a Mitacs award (BW), the European Union’s Horizon 2020 research and innovation program under a Marie Sklodowska-Curie grant (ER), by the CIFAR Azrieli Global Scholar program (JPJ), by the Ontario Early Researcher Awards program (JPJ and CNA), and by the Canada Research Chairs program (JPJ). We also thank Dr. James Rini (University of Toronto) for the kind gift of the 9.8E12 antibody used to detect the 229E Spike protein, and Dr. Scott Gray-Owen (University of Toronto) for the kind gift of the NL63 human coronavirus.

## Supplemental Materials

**Table S1.** Cell culture reagents for propagation of human coronaviruses

| **Name** | **Component** | **Company** | **Catalogue #** | **Final Amount** |
| --- | --- | --- | --- | --- |
| **Growth Media** | Minimum Essential Media Eagle | Millipore Sigma | M4655 | N/A |
|  | Fetal Bovine Serum, Heat-inactivated | Thermo Fisher | 10082147 | 10% |
|  | Penicillin-Streptomycin | Thermo Fisher | 15070-063 | 1X |
| **Infection Media** | Minimum Essential Media Eagle | Millipore Sigma | M4655 | N/A |
|  | Fetal Bovine Serum, Heat-inactivated | Thermo Fisher | 10082147 | 2% |
|  | Penicillin-Streptomycin | Thermo Fisher | 15070-063 | 1X |
| **Plaque Media** | MEM (Temin's modification) (2X), no phenol red | Thermo Fisher | 11935046 | 1X |
|  | Fetal Bovine Serum, Heat-inactivated | Thermo Fisher | 10082147 | 1% |
|  | Penicillin-Streptomycin | Thermo Fisher | 15070-063 | 1X |
|  | 0.6% Agarose | Thermo Fisher | 16500-500 | 0.3% |
| **Transfection Master Mix** | Opti-MEM- I Reduced-Serum Medium | Thermo Fisher | 31985070 | N/A |
| | Lipofectamine RNAiMAX | Thermo Fisher | 13778030 | 6.3 μL per well |
|  | siRNA (control or target) | N/A | N/A | 50nM |

**Table S2.**
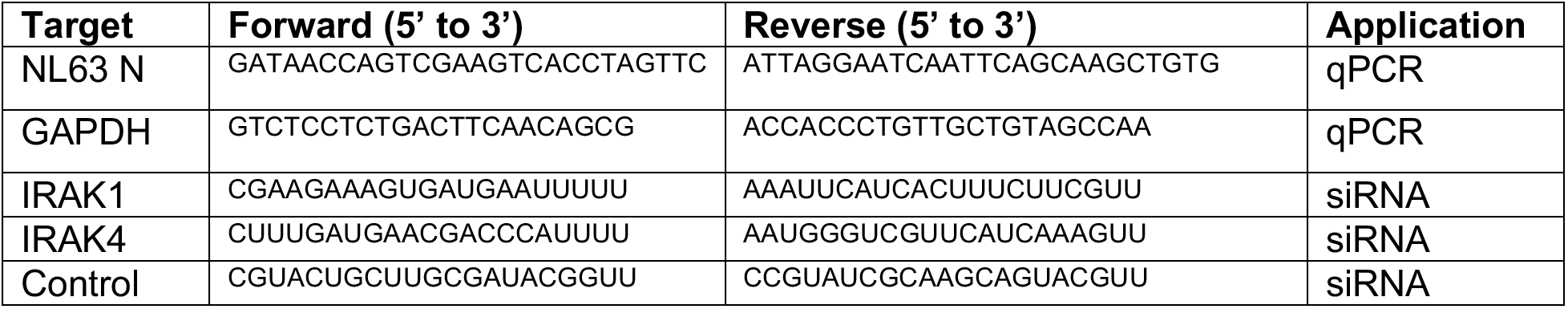
Oligonucleotide sequences for siRNA transfection or qPCR

**Table S3.** RT-qPCR protocol

| **Step** | **Temperature** | **Time** |
| --- | --- | --- |
| 1. Reverse transcription | 55°C | 10 minutes |
| 2. Hot start | 95°C | 1 minute |
| 3. Denaturation | 55°C | 10 seconds |
| 4. Extension (with plate read) | 62°C | 30 seconds |
| 5. Repeat 3-5 (45x) | N/A | N/A |
| 6. Melt curve | 60-95°C, 0.5°C increments | 5 seconds |

**Table S4.** Additional information for compounds used in this study

| Number | Compound | Supplier | Cat. No. | Final Concentration |
| --- | --- | --- | --- | --- |
| 1 | Bucladesine | MedChemExpress | HY-B0764 | 10 μM |
| 2 | Cinnarizine | MedChemExpress | HY-B1090 | 1 μM |
| 3 | Doxycycline | MedChemExpress | HY-N0656 | 10 μM |
| 4 | Eflornithine | MedChemExpress | HY-B0744 | 10 μM |
| 5 | Flucytosine | MedChemExpress | HY-B0139 | 10 μM |
| 6 | Opicapone | MedChemExpress | HY-14896 | 1 μM |
| 7 | Otilonium Bromide | MedChemExpress | HY-B0499A | 1 μM |
| 8 | Cefotiam | MedChemExpress | HY-B0734A | 1 μM |
| 9 | Dapagliflozin | MedChemExpress | HY-10450 | 1 μM |
| 10 | Fosamprenavir | MedChemExpress | HY-78726 | 10 μM |
| 11 | Gentamicin | MedChemExpress | HY-A0276 | 10 μM |
| 12 | Glipizide | MedChemExpress | HY-B0254 | 10 μM |
| 13 | Palbociclib | MedChemExpress | HY-50767 | 1 μM |
| 14 | Saquinavir | MedChemExpress | HY-17007 | 1 μM |
| 15 | Streptozotocin | MedChemExpress | HY-13753 | 10 μM |
| 16 | Sugammadex | MedChemExpress | HY-B0079 | 10 μM |
| 17 | Tofacitinib | MedChemExpress | HY-40354 | 10 μM |
| 18 | Capmatinib | MedChemExpress | HY-13404 | 10 μM |
| 19 | Anidulafungin | MedChemExpress | HY-13553 | 1 μM |
| 20 | Glibenclamide | MedChemExpress | HY-15206 | 10 μM |
| 21 | Nelarabine | MedChemExpress | HY-13701 | 1 μM |
| 22 | Remdesivir | MedChemExpress | HY-104077 | 4 μM |
| 23 | Degarelix | MedChemExpress | HY-16168A | 1 μM |
| 24 | JH-I-25 | Millipore-Sigma | SML2609 | 10 μM |
| 25 | SP600125 | Millipore-Sigma | S5567 | 10 μM |
| 26 | SB202190 | abcam | ab120638 | 10 μM |
| 27 | PD98059 | abcam | ab120234 | 10 μM |

**Figure S1.**
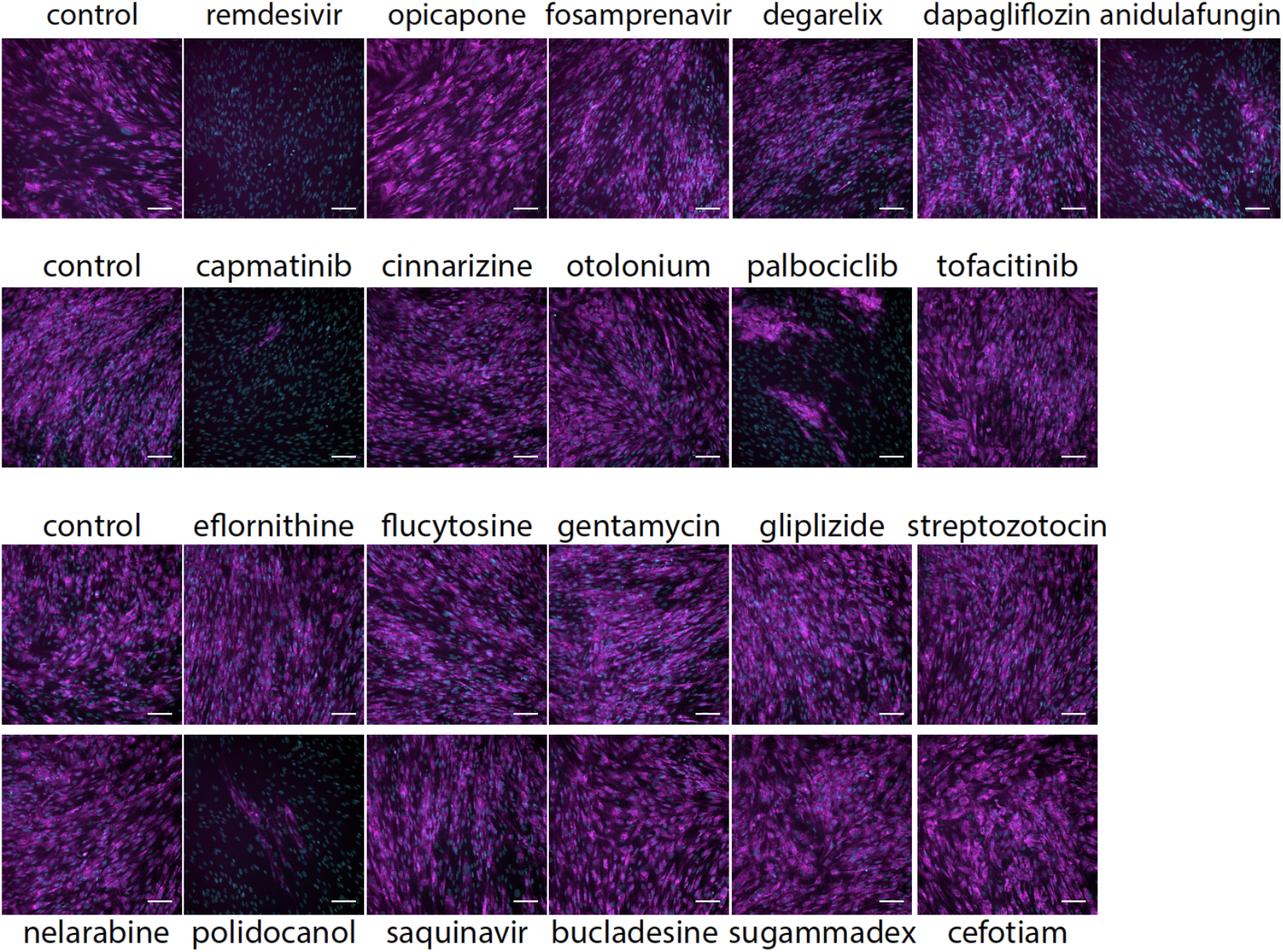
Representative microscopy images for data shown in Figure 2. Representative images from MRC-5 cells treated with drugs as indicated (see **Table S4**) and infected with 229E for 2 days. Images depict S protein expression (magenta) or DAPI (cyan) Scale, 100 μm.

**Figure S2.**
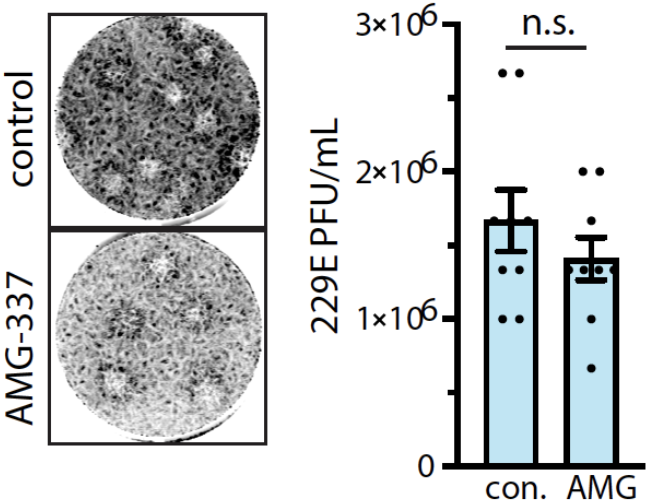
Inhibition of MET by AMG337 does not impact 299E infection assessed by PFU assay. (left) Representative images of plaques in MRC-5 cells treated with DMSO (vehicle) control or 10 μM AMG-337 and infected with 299E and (right) quantification of 299E viral titer shown as mean PFU ± SE (n = 3 with 3 technical replicates per experiment).

**Figure S3.**
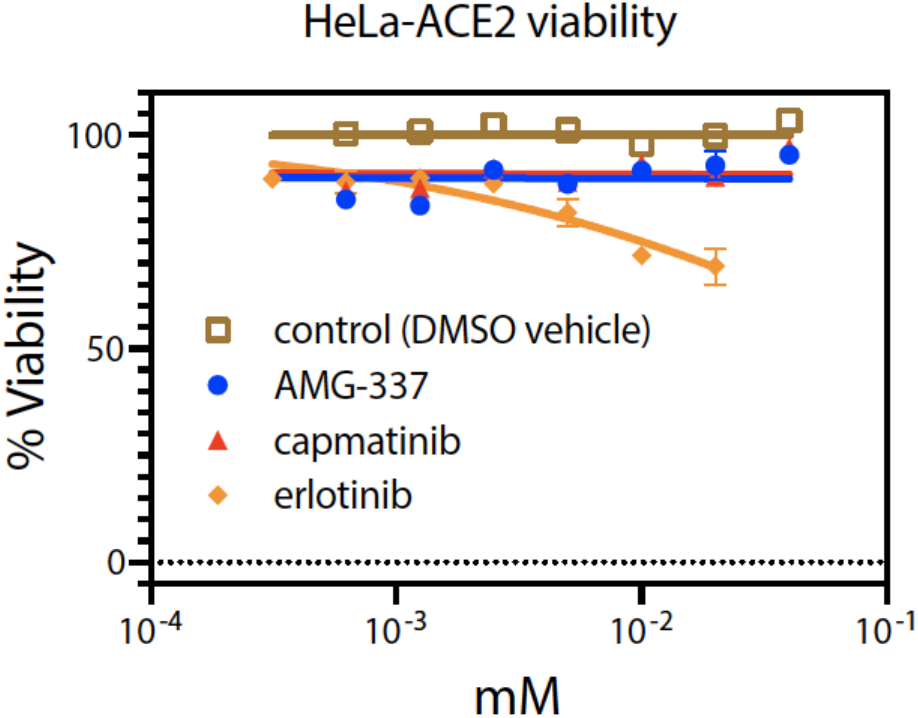
Capmatinib does not affect cell viability. HeLa-ACE2 cells were treated with inhibitors as shown (10 μM each) in parallel with the PsV assay shown in Figure 4D. Shown are representative cell viability measurements from n=3 independent experiments. Neither capmatinib nor AMG-337 impact cell viability. In contrast, the EGF Receptor inhibitor erlotinib resulted in a dose-dependent impairment of cell viability.

**Figure S4.**
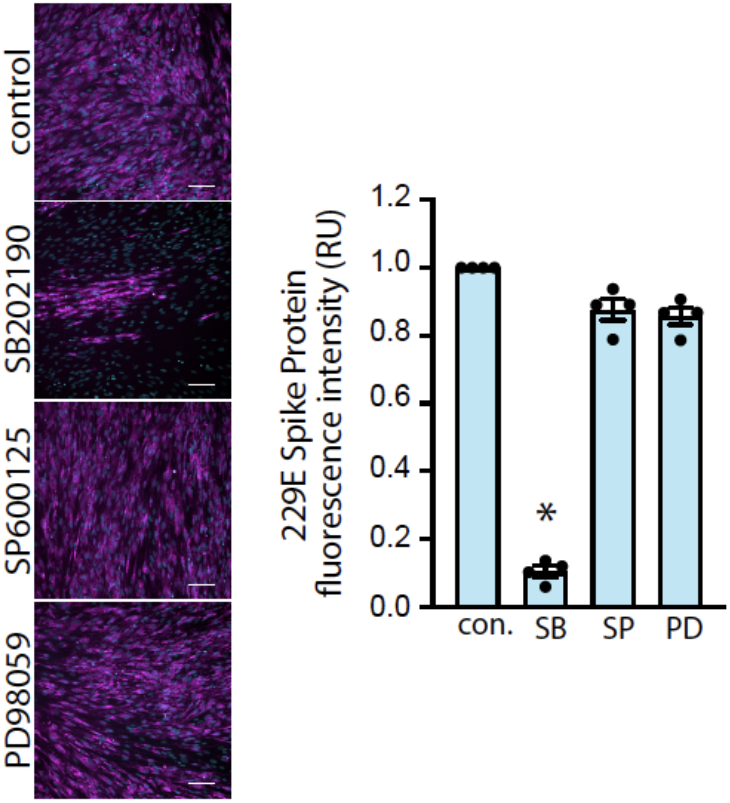
Inhibition of the p38 MAPK recapitulates the effects of capmatinib and IRAK1/4 inhibition on coronavirus infection. (left) Representative images from MRC-5 cells treated with 10 μM SB202190 (SB, p38 MAPK inhibitor), 10 μM SP600125 (SP, JNK inhibitor) or 10 μM PD98059 (PD, MEK1 inhibitor) and infected with 229E for 2 days. (right) Quantification of 229E S protein expression (>10 images per condition, n = 4).

